# Neural RNA splicing models rely on reading frame information unavailable to the splicing machinery

**DOI:** 10.64898/2026.11.03.763976

**Authors:** Kavi Gupta, Bixuan Wang, Phillip A. Sharp, Christopher B. Burge, Armando Solar-Lezama

## Abstract

Deep neural networks can accurately predict RNA splice sites from genomic sequence, but whether they do so by simulating the natural splicing process is unclear. We evaluate this question by examining mutations that should not have a strong effect on splicing, but are correlated with it in the human genome. We focus specifically on mutations that affect frame alignment because this is known to be relevant to machinery that operates spatially and temporally separately from the splicing machinery. We find that predictions made by commonly-used neural tools are impacted far more by deletions or mutations that alter translational reading frame, producing in-frame stop codons, than by those that preserve reading frame. This dependence on reading frame causes these models to systematically miss many poison exons, a clinically relevant category of exons where reading frame is disrupted. This dependence is much weaker in “grey-box” models that build in splicing-specific structure. Our results imply that widely-used black-box models of splicing rely on information that is not accessible to the RNA splicing machinery. Code is available at github.com/kavigupta/frame-alignment-checks.

## 1 Introduction

The expression of most human genes requires the excision of multiple introns from the primary RNA transcript by the splicing machinery to produce mature mRNA that will direct protein synthesis. The sequence features that underlie the choice of splice sites by the splicing machinery include core splice site motifs and diverse splicing regulatory elements (SREs), whose functions are only partially understood. There is a growing literature on the use of machine learning to identify SREs and infer their behavior. Some of these methods [Gupta et al., 2024, McCue and Burge, 2024] aim to explicitly capture how splice site motifs and SREs influence splicing. However, the most accurate prediction of splice site locations on human genes has been achieved by black-box models such as SpliceAI, Pangolin and others trained on thousands of annotated human genes, which have greater than 90% accuracy [Jaganathan et al., 2019, Zeng and Li, 2022].

These models have a generic structure and thus rely on information gained in training rather than domain-specific prior knowledge. This approach allows the models to potentially discover signals that are unknown in prior work, but it also introduces the risk that the models can learn to rely on signals that are different from those used by the actual biochemical splicing machinery. This is a general risk with any application of machine learning, and genomic sequence data is especially likely to expose this vulnerability. Gene expression involves several different steps that use distinct mechanisms that rely on distinct control signals layered in the same underlying sequence. These signals can be highly correlated even when they are not causally linked. A model that learns to rely on these correlated signals may achieve high accuracy, but the learned function may not end up corresponding to any specific biological mechanism. We have concrete evidence that despite high accuracy, black-box models perform poorly under distributional shift to non-genomic sequences (Appendix A).

In this paper, we demonstrate that some of this poor performance can be explained by models learning a non-causal correlation. Specifically, multiple state-of-the-art neural splicing predictors exploit protein-coding information that is not part of the biochemical splicing code. This leads to systematic prediction errors for exons that do not form part of the canonical translation reading frame, such as *poison exons* (alternatively spliced exons that carry an in-frame stop codon) and exons in non-coding RNA genes. The protein translation machinery expects an open reading frame (ORF) in order to be able to translate mRNA into a protein. This feature (see Section 2.4 for details) only has about a 38% chance of appearing in a random sequence of 120nt but must appear in nearly every coding exon. As a result, the ORF signal is highly correlated with splicing, despite being causally independent. For example, the splicing machinery includes poison exons in many genes, which are often involved in gene expression regulation [Ni et al., 2007], but do not necessarily contain an ORF.

We introduce a battery of in-silico mutagenesis tests that can quantify the use of reading frame information by a given splicing model, and detect the ORF reliance mentioned above. To confirm that the splicing machinery does not in fact use this signal, we analyze results of experimental splicing reporter assays and poison exons in human tissues. We find that models that more strongly leverage ORF information in the in-silico experiments underpredict exons without an ORF compared to experimentally measured values, confirming that this dependence is not reflected in actual splicing outcomes. We discuss the implications of these observations for the design of datasets and model architectures for learning mechanistic rules of gene expression.

## 2 Preliminaries

### 2.1 RNA Splicing

Cells use a series of complex procedures to transform information stored in DNA into usable proteins. One step along the way involves taking pre-mRNA (Figure 1a), which represents the contents of a gene, and removing several *introns*, to produce the mRNA (Figure 1c). The sequences within the pre-mRNA that are not removed are called *exons*. This includes the portion before the first intron and after the last intron, but in this work we focus on *internal exons*, the ones that appear between introns.

**Figure 1:**
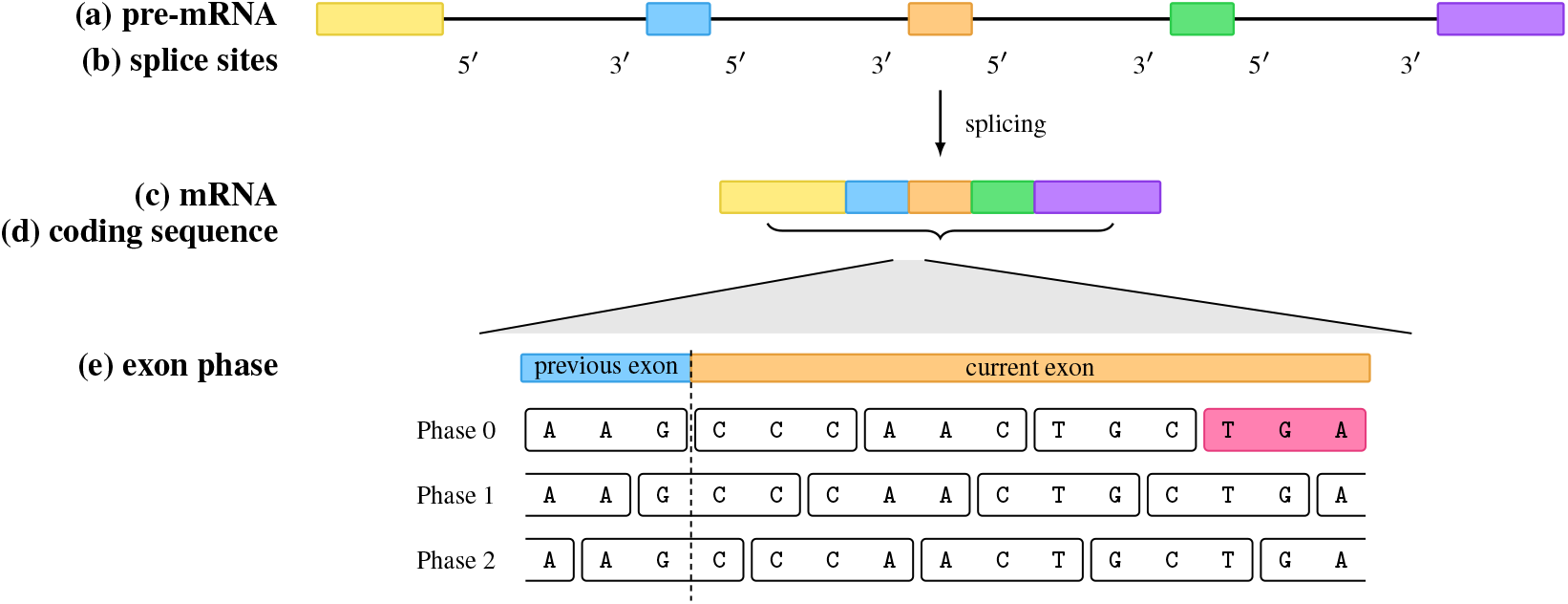
**(a)** A pre-mRNA, with exons shown as colored blocks and introns as the line in between them. **(b)** Every internal exon begins at a 3^/^ splice site (3SS) and ends at a 5^/^ splice site (5SS). **(c)** Splicing out the introns results in the mRNA, composed of exons. **(d)** The coding sequence is a subsequence of the mRNA. **(e)** Zooming in on an exon–exon junction of the mRNA. Three possible views of the codon tiling depending on how the previous exon’s tiling ended. The phase is the position of the start of the exon relative to the coding sequence. In this case, Phase 0 has an in-frame stop, whereas Phases 1 and 2 do not.

Some internal exons are *constitutive*, appearing in essentially every mRNA produced from the gene, while others are *alternative*, included only in some transcripts; the fraction of transcripts including a given exon is its *splicing level*.

### 2.2 Models

All splicing models we discuss focus on the problem of identifying *splice sites* (Figure 1b), that is, the boundaries between exons and introns. Whenever we report the accuracy of an exon prediction, it is the average accuracy in predicting the start (3’ splice site) and end (5’ splice site) of the exon.

We analyze both *black-box* and *grey-box* models. Both categories of models are trained on large datasets of human genomic sequence annotated with splice sites, but whereas the black-box models we analyze are generic deep neural networks, the grey-box models we analyze produce intermediate predictions of binding sites of RNA binding protein (RBP) motifs, and exclusively use this information to predict splice sites. This constrains them to be more mechanistic, and less able to exploit correlations in the training data not causally related to splicing, but also makes them less accurate.

We analyze the following black-box models: SpliceAI (400nt and 10knt window variants) [Jaganathan et al., 2019], Pangolin [Zeng and Li, 2022], and AlphaGenome [Avsec et al., 2026]. We analyze the grey-box models SAM-FM (fixed motifs) and SAM-AM (adjusted motifs) [Gupta et al., 2024], where SAM-FM is a model that uses a neural network to predict splice sites from predicted binding of RNA binding proteins using known binding motifs, and SAM-AM is a model that adjusts the binding motifs using end-to-end learning signal. Of note is that SpliceAI, Pangolin, and SAM are trained on data processed in a roughly similar way to remove extremely rare variants (SpliceAI and SAM are trained on near-identical training sets, while Pangolin does incorporate more information about tissue-specific and alternative splicing); on the other hand, AlphaGenome is a more recent model that trains on nearly unfiltered exons.

### 2.3 RNA Translation

Within the mRNA, a certain portion of the text is the *coding sequence* (CDS, Figure 1d), which is used to code for proteins. This sequence begins with an ATG^2^ and is read in groups of 3nt called *codons*, ending with a *stop codon*, which is one of TAG, TAA, and TGA. In general, stop codons cannot appear in this codon sequence in internal exons, except within the final 50nt of the last internal exon; if they do, this is a premature stop and leads to the transcript being degraded by Nonsense-mediated mRNA Decay (NMD) [Margasyuk et al., 2024]. This means that evolution constrains nearly all internal exons to have a tiling of codons where none are stops [Rong et al., 2020].

### 2.4 Open Reading Frames

As seen in Figure 1e, an exon can be in one of three *phases* relative to the codon sequence. Where the codon begins depends on where in the coding sequence the previous exon ended. In practice, only one of these will ever be read, but this is an exclusively global property. Looking only at one exon, all we can say is that at least one of these 3 phases must not contain a stop codon. A sequence has an *open reading frame* (ORF) if there exists a phase in which no stop codon appears in the codon sequence.

In contrast, introns have no such requirement. A typical exon is about 120nt (40 codons) long. Since 3 of 64 RNA triplets represent stop codons, the probability that a randomly generated sequence of length *L* nucleotides will have an ORF is approximately 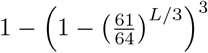. Thus, a random segment of length 120 only has a~ 38% chance of having an ORF. This makes the presence of an ORF a powerful signal for splice site prediction, even though the splicing machinery does not depend on frame information.

Liu et al. [2026] note that splicing predictors are often highly sensitive to stop codons, and that this sensitivity is 3-periodic, suggesting a connection to codons. However, they do not identify the specific property that models are picking up on: the open reading frame. This difference is important as stop codons are often splicing features (as many other triplets are), whereas the presence of an open reading frame is not accessible to the splicing machinery.

## 3 In-Silico Mutagenesis

In order to measure the degree to which models are using ORF signals, we performed several large-scale in-silico mutagenesis experiments. In each experiment, we made isolated mutations to annotated exons that are expected to typically have limited effects on splicing, and measured changes in the accuracy of the models in predicting the exon’s splice sites. Specifically, we analyze frame-disrupting deletions and substitution of sense codons by stop codons.

### 3.1 Methodology

In all these experiments, we start with the first half of the genes in the SpliceAI test set (what Gupta et al. [2024] use as the validation set) containing 831 genes. Of these, 635 satisfy the conditions: (1) appear in the Ensembl annotation database [Harrison et al., 2024], (2) have exactly one transcript matching the canonical transcript SpliceAI uses in its dataset, (3) have an annotated coding sequence (CDS), (4) have at least one internal exon, (5) have a start codon, and (6) have no in-frame stop in the CDS. We then produce a list of 4954 internal coding exons from these 635 genes. Finally, we filter these exons for length at least 100nt, leaving us with 3156 exons.

Though all the models we analyze produce values in the output range (0, 1), there is no reason to specifically treat these as grounded probabilities or on the same scale as each other. As a result, we use these models as binary predictors, setting the threshold to the top-k threshold (the threshold such that the number of false positives and false negatives on the SpliceAI test set is identical). We then report changes in accuracy, as if the mutation we are performing does not change ground truth (i.e., a drop in accuracy means some of our exons stop being predicted as such). See Section 4 for justification of this assumption.

### 3.2 In-silico deletion

First, we studied deletions of 1 to 9 nucleotides (nt). Exonic deletions of 3, 6 or 9nt preserve the exon’s phase (and thus the reading frame), while other deletions are phase-altering. Intronic deletions and phase-preserving exonic deletions serve as controls for the phase-disrupting exonic deletions. The deletions were located 40nt upstream or downstream from the nearest 3SS or 5SS (Figure 2a). As seen in Figure 2b, intronic deletions had minimal effects on predictions for all models. However, exonic deletions had larger effects, with the proximal SS impacted more strongly than the distal SS. Strikingly, a strong period-3 pattern was observed for the SpliceAI and Pangolin models, with frame-disrupting deletions reducing accuracy by 10–20% in most cases, while frame-preserving deletions had negligible effects on accuracy. A more modest effect in the same direction was observed for the grey-box SAM-AM model, and an even more modest effect with SAM-FM [Gupta et al., 2024]. These observations imply that black-box models of splicing are strongly sensitive to disruption of reading frame and/or to the disruption of 3-periodic patterns in coding exons, as we had hypothesized. Interestingly, AlphaGenome also had a fairly minimal effect, which we hypothesize is due to its much larger and less well-filtered training set (Section 2.2) avoiding the bias we explore here. In the remainder of this paper, we focus on the other black-box models, as they remain in common use, and the dataset bias issue might re-occur in future models that train on more curated datasets.

**Figure 2:**
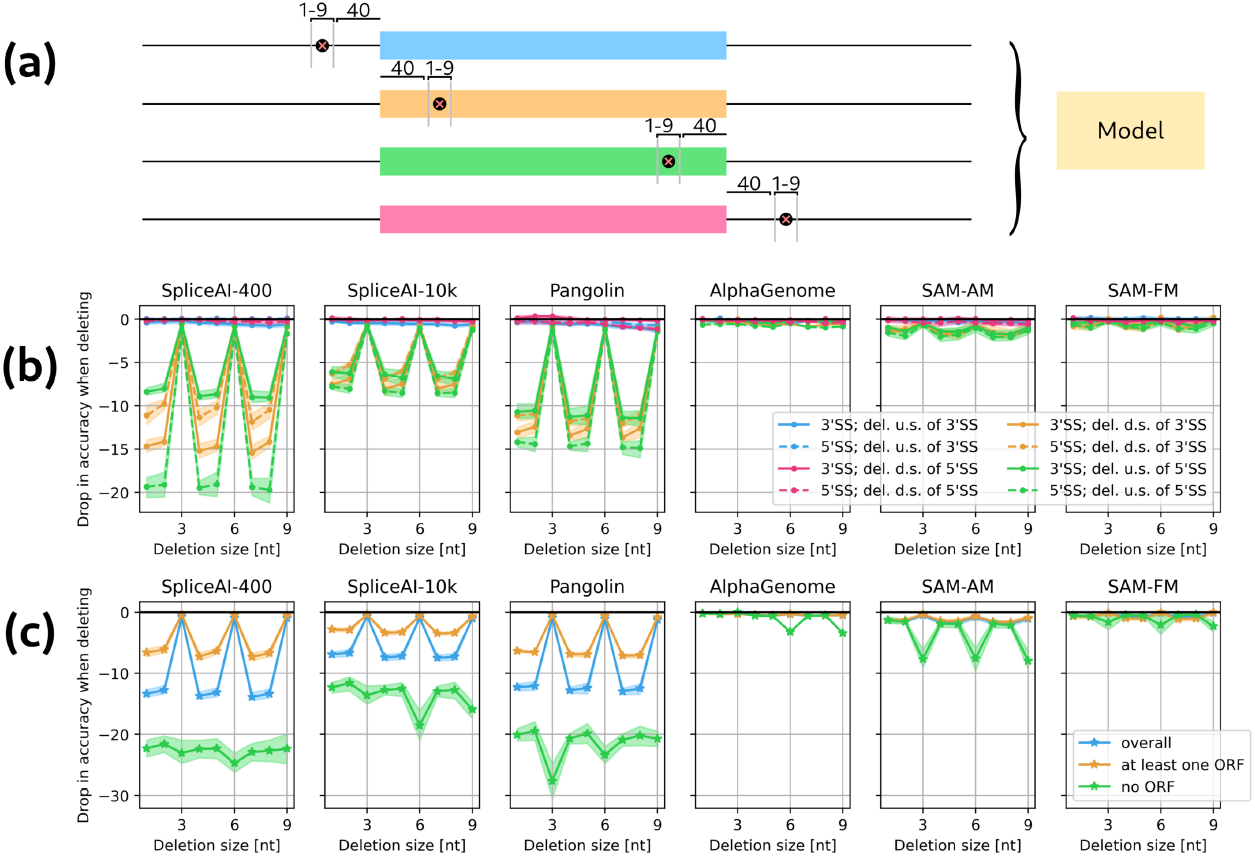
In-silico deletion experiment. **(a)** Diagram of in-silico deletion experiment, in which a short segment of 1 to 9nt is deleted at a position 40nt upstream or downstream of the 3SS or 5SS of an exon. **(b)** The drop in accuracy of 3SS and 5SS prediction is shown for each of 4 types of deletion for 6 splicing models (del.: deletion; u.s./d.s.: upstream/downstream). In each case, the mean over 5 replicate training runs is shown (except for AlphaGenome, which is the result of querying the API), with shaded regions indicating 95% bootstrap confidence intervals. Intronic deletions and frame-preserving exonic deletions have minimal effects on accuracy, while exonic deletions that alter frame substantially reduce accuracy of both splice sites, especially for SpliceAI and Pangolin. **(c)** The impact on SS prediction accuracy of short deletions grouped by whether or not they result in every one of the exon’s reading frames being closed.

### 3.3 Models seek an Open Reading Frame

We hypothesized that these models might have learned to distinguish inauthentic “pseudoexons” (decoy sequences that resemble exons but are not spliced in) by the absence of a gene-spanning open reading frame. However, we observed that deletions in one exon minimally impacted prediction of splice sites of adjacent exons (Appendix C.1), arguing against detection of a gene-spanning ORF feature. Furthermore, frame-altering mutations in one exon reduced accuracy, even when they were compensated by an offsetting deletion in an adjacent exon that repaired reading frame (Appendix C.2). Therefore, it appears that these models identify ORFs in individual exons and not across multiple exons.

The remaining relevant feature specific to individual exons is whether the exon has an ORF in any of the possible frames. To investigate this feature, we reanalyzed the deletions from Figure 2b by whether the resulting exon contained or lacked any ORF. As seen in Figure 2c, deletions that disrupted all ORFs caused much greater drops in accuracy compared to those that left a reading frame open. Furthermore, in the rare case when a 3, 6 or 9nt deletion introduced an in-frame stop codon at the deletion site, this also lowered accuracy substantially. These observations confirm that these models have learned to detect presence/absence of an ORF in putative exons. However, frame-altering deletions still reduced accuracy relative to frame-preserving deletions, even when an ORF remained in the exon. This suggests that these models are also recognizing other coding-related information, for example period-3 patterns arising from biases in amino acid and codon usage [Staden, 1984, Fickett, 1982, Wang and Stein, 2010].

### 3.4 In-silico replacement of 3mers with Stop Codons

Previously, it has been observed that mutations which create stop codons often disrupt positive-acting exonic splicing enhancer (ESE) motifs and/or create negative-acting exonic splicing silencer (ESS) motifs [Rong et al., 2020, Cartegni et al., 2002]. To further dissect the effects of exonic sequence on predictions, we performed all 64 possible 3nt substitutions^3^ in each of the three possible reading frames~ 40nt away from splice sites in our set of 3,156 exons (Figure 3a). Like in the deletion experiment, we look at locations 40nt away from the splice sites and then move downstream 0, 1, or 2nt until we are in the appropriate phase relative to the gene-level coding sequence.^4^

**Figure 3:**
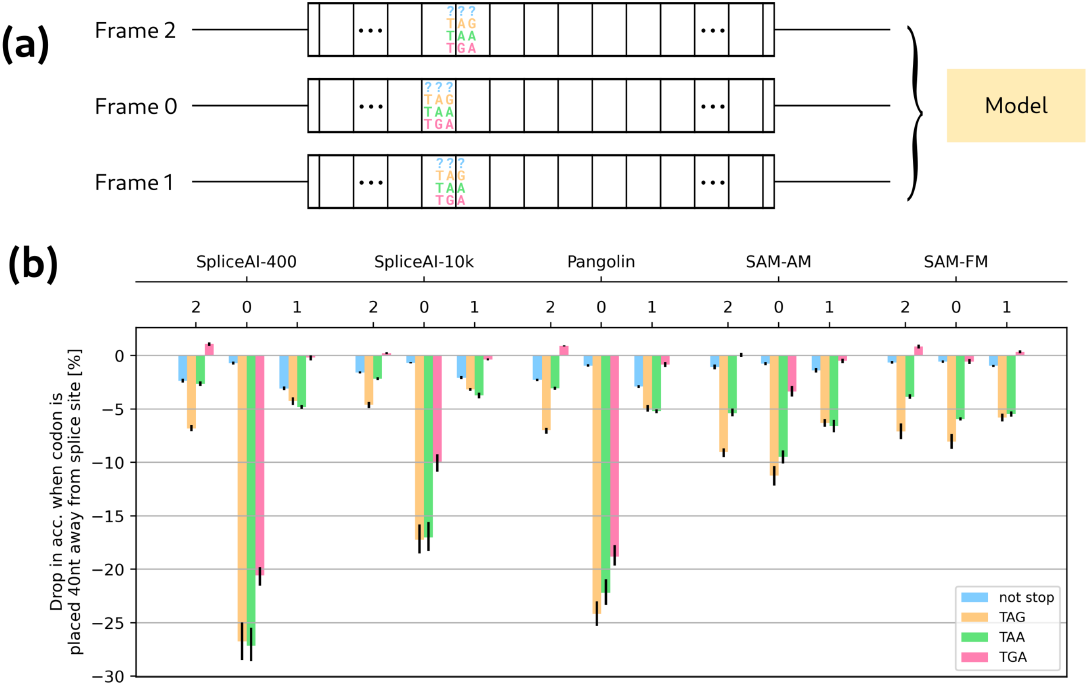
In-silico 3mer replacement experiment. **(a)** Diagram of an in-silico mutagenesis experiment, in which an individual RNA triplet was replaced by another triplet, at one of four locations~ 40nt downstream of the 3’ splice site (as shown) or upstream of the 5’ site of an exon, either in-frame or in the 2 or 1 reading frame. Mutations that created or removed a stop in a frame other than the one they were inserted into (e.g., `CCCAG` → `TTTAG`) were excluded. **(b)** The impact of triplet substitutions on SS prediction accuracy. SpliceAI and Pangolin are highly sensitive to substitution of a codon by a stop codon, but not by similar substitutions out-of-frame.The impact on SAM models depends on which stop codon triplet is inserted, but SAM models are much less sensitive to in-frame versus out-of-frame stop codon substitutions. Results for all individual codons and frames are discussed in Appendix B.

In general, introduction of in-frame stop codons caused larger reductions in accuracy than introduction of other codons (Figure 3b). Furthermore, this effect is primarily present when stop codons are placed in-frame, with out-of-frame stop codon triplets having much smaller effects. In fact, placement of stop codons in-frame caused larger drops than any of the 189 other substitutions considered (see Appendix B for more details). The relative impact of in-frame stops on accuracy indicates that SpliceAI and Pangolin rely heavily on reading frame information, while SAM-AM uses it modestly, and SAM-FM uses it minimally.

Introduction of out-of-frame TAA and especially TAG triplets also reduced accuracy, while TGA triplets were largely neutral or even slightly positive in some models (Figure 3b). These latter observations can be explained in terms of known splicing regulatory activities of motifs containing these triplets that these algorithms may have learned to some extent. For example, TAA and especially TAG were highly enriched in a cell-based screen for ESSs [Wang et al., 2004], but were absent from a standard set of ESEs [Fairbrother et al., 2002], while TGA occurred many times in the ESE set and only once in the ESS set. Thus, the effect of the TGA triplet provides the most direct measure of the extent to which an algorithm relies on reading frame information. Together, these observations indicate that black-box splicing models are specifically sensitive to stop codons when they are present in-frame.

## 4 Empirical evidence confirms frame signal is non-mechanistic to splicing

One potential objection to the results in Section 3 is that, while it is generally believed that reading frame does not impact splicing, this might be incorrect, and splicing models have simply picked up on this true signal. In this section, we demonstrate that these models’ dependence on reading frame is a bias relative to *ground truth* changes in splicing when reading frame is disrupted.

### 4.1 Controlling for true splicing

In order to assess this claim, we need to be able to control for true splicing. To accomplish this, we use datasets of experimentally verified splicing measurements on perturbed exons and collect for each exon three pieces of information: a proxy for true splice site strength, the model’s prediction, and whether it has an ORF. This allows us to use the ORF-containing exons as controls for the no-ORF exons.

As Figure 4 shows, we compared the scores assigned to exons^5^ lacking an ORF to the scores of control exons with reading frame(s) open, choosing as controls the 100 exons which had the most similar experimentally-measured splicing levels. We then tallied the proportion (out of 100) of these control exons with scores above the score for the target exon, calling this proportion “fraction of predictions above” (FPA).

**Figure 4:**
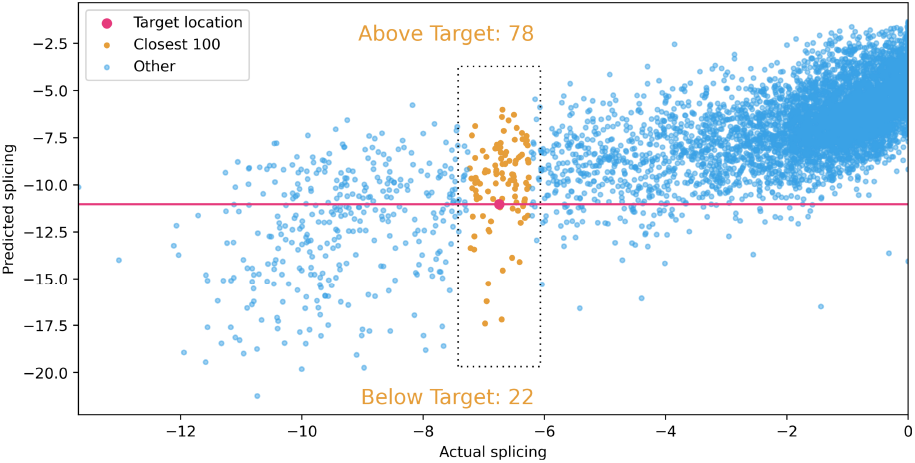
Illustration of the FPA (fraction of predictions above) calculation. For a given target exon with all reading frames closed (red dot), we calculate the scores of the 100 control exons with the most similar measured splicing levels (orange dots), and ask what fraction of these control exons have scores above that of the target exon.

The intuition is that if a feature is being appropriately scored then exons with that feature should have scores similar to other exons with similar splicing levels, hence FPA near 50%. On the other hand, if exons with no ORF are scored substantially below the median of other exons with similar splicing levels, this would indicate that such exons are underpredicted relative to their true splicing level.

### 4.2 Datasets

We use two kinds of datasets here. In both cases, we minimize the role of NMD by ensuring that either it affects none of the exons or all of them.

#### Minigenes

We searched the literature for experimental measurements of the splicing impacts of large sets of mutations or deletions similar to those we generated in-silico, finding two suitable datasets. Both come from *minigene* reporters: short artificial genes carrying an exon of interest. One recent study (referred to as “Saturation Mutagenesis” here) analyzed the impact of over five thousand 2nt and 6nt replacement mutations on splicing of a 3-exon minigene transfected into human cells [Ke et al., 2018]. Another recent study (referred to as “Deep Indel”) analyzed the impacts of over five thousand small insertions and deletions on splicing [Baeza-Centurión et al., 2025]. In the Saturation Mutagenesis experiment, 77 exons lacked an ORF, and 5483 had at least one frame open. Meanwhile, the Deep Indel dataset yielded 282 exons with no ORF and 5695 with at least one frame open. For ground truth, we used the splicing level estimates reported by the authors in the corresponding studies: log_2_(EI) and log_2_(PSI),^6^ respectively.

Both studies used a similar splicing reporter system with no start codon, minimizing the role of NMD. In the analysis of these data, we focused on the central question of whether splicing models systematically underpredict splicing for exons that lack an ORF relative to other exons.

#### Poison Exons

Underprediction of the splicing of potential exons lacking an ORF likely improves accuracy in the prediction of annotated coding exons, but may impede prediction of alternative splicing, which often alters reading frame [Lewis et al., 2003]. “Poison cassette” exons are internal, alternatively spliced exons that contain a premature stop codon in-frame relative to upstream exons. Such exons are common in mammalian genomes and can play important roles in regulating gene expression, but are absent from many standard annotations, including those used to train and test the splicing models studied here [Carvill and Mefford, 2020]. Manipulation of the splicing of such exons can be used to up- or down-regulate gene expression for therapeutic purposes, drawing significant interest from biopharma [Bakshi and Isom, 2025].

Poison exons can be challenging to detect from transcriptomic data because isoforms that contain them are often efficiently degraded by NMD [Margasyuk et al., 2024]. For the same reason, their PSI levels as measured in standard RNA-seq may substantially underestimate the levels at which they are spliced in the nucleus. We wondered whether these models have effectively learned the aggregate of splicing+NMD, rather than purely splicing. To study human poison exons at scale, we used a large dataset of “Naturally Variable Exons” (NVEs), defined as exons which are included in some but not all members of a population; they are largely unannotated and often disrupt reading frame [Jacobs et al., 2025]. NVEs have been systematically identified using the GTEx dataset of tissue RNA-seq samples from over 800 individuals [Lonsdale et al., 2013]. We filtered for poison NVEs by requiring that they contain an in-frame stop (relative to their surrounding exons) that is at least 50nt upstream of the last intron in the gene. To control for the effects of NMD, we focused on comparing poison exons that have a stop codon in all 3 frames to poison exons where at least one of the other two frames is open. Since both sets contain stop codons in frame with upstream exons, they should be equally sensitive to NMD. For ground truth, we used “exon frequency” at 5% (EF 5%, the fraction of individuals that include the exon with PSI at least 5%), which is a robust measure that is highly correlated with PSI.

### 4.3 Results

Examining the mean FPA for all exons lacking ORF (Figure 5), we find that, across all datasets, black-box models (SpliceAI, Pangolin) predict lower scores for these exons than their splicing levels merit, while this effect is only observed in one dataset for grey-box models (SAM-FM, SAM-AM). Importantly, this analysis uses measured splicing levels to control for any effect that the presence of stop codons might have on splicing per se. Varying the number of control exons did not substantially affect results (see Appendix D.2). The relationship between measured splicing level and predicted score across exons is shown in Figure 12. We observed an analogous trend when using a variant of this metric where the splicing levels of exons lacking an ORF are compared to those of control exons with similar scores (Appendix D.1).

**Figure 5:**
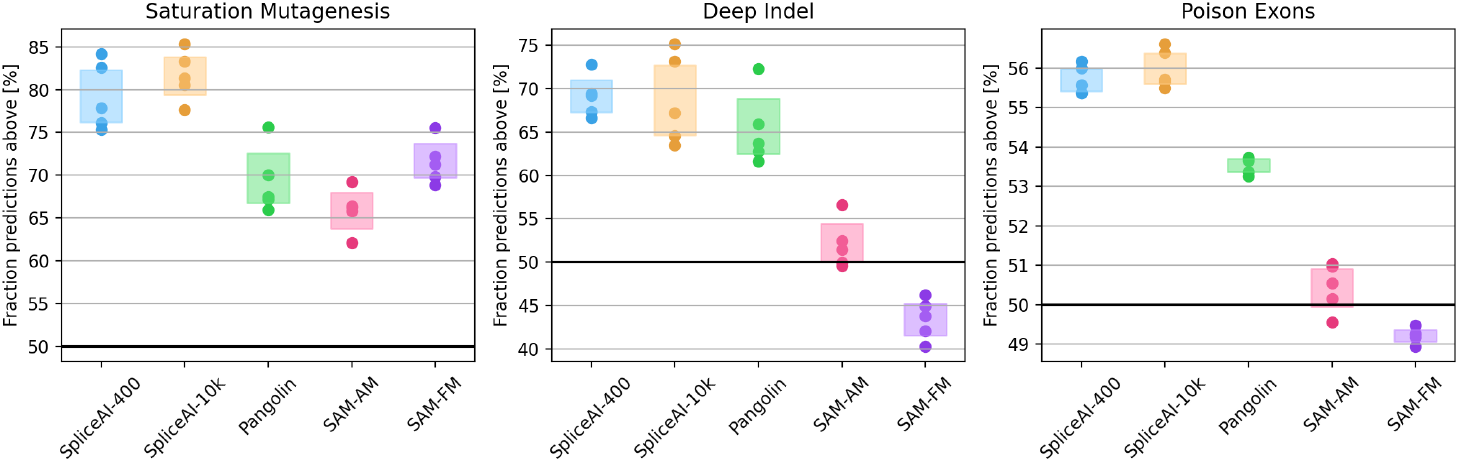
FPA results on minigene data and poison exons. Each point represents a replicate training run, and boxes represent a 95% bootstrap CI. The black line marks FPA of 50%, the value expected if models were not underpredicting exons without an ORF; values above 50% indicate model underprediction. More detailed plots are shown in Appendix E.

The results on poison exons demonstrate that black-box models significantly underpredict poison exons that lack any ORF relative to other poison exons, despite both groups being equally subject to NMD. Thus, the models’ preference for ORFs cannot solely be a reflection of them having learned the rules of NMD. They have developed an artificial preference for potential exons which have at least one reading frame open relative to those that have a stop codon in all three frames. This property is useful in prediction of coding exons, but does not reflect a natural biochemical mechanism since exons are typically translated in just one frame.

## 5 Discussion

Together, the results here demonstrate that widely-used black-box models of splicing closely resemble gene-finding algorithms [Burge and Karlin, 1998], which use protein-coding information in addition to splicing information to identify coding exons.

This conclusion has several implications. One is that black-box models should not be relied on to understand the mechanism of splicing, since they rely on types of information not accessible to the splicing machinery. A second implication is that, in applications such as variant interpretation, black-box model predictions for variants that alter reading frame are likely to substantially overestimate their true impact on splicing. Another implication is that in applications to identify alternative exons, black-box models will tend to underpredict or completely miss poison exons, an important class for potential therapeutic manipulation of gene expression.

Gene function is a multi-step process where sequence features relevant to all steps are interleaved and evolve jointly. In this paper, we explored how one particular pair of correlated features impacted the ability of black-box models to learn the splicing mechanism, but this problem could apply to any mechanism and any features. Thus, black-box models that seek to mimic particular steps in gene expression will tend to identify correlated rather than causal information, unless very strict guardrails are erected during design and training to enforce or at least enrich for causal features. These observations suggest that alternative approaches, such as the use of grey-box models, are needed to more faithfully model the underlying processes of gene function.

## Acknowledgments and Disclosure of Funding

We thank Juan Valcárcel for providing helpful feedback on an early manuscript draft. This work was supported by the National Science Foundation under Grant No. CCF-1918839 and by MIT HEALS.

## A Evaluation of Models on Non-Genomic Datasets

We evaluate a suite of models trained on the genome on two non-genomic datasets, demonstrating significant limitations in the generalizability of these models. These sequences are not under the evolutionary constraint of being translated to proteins that keeps a frame open in real coding exons, so some of the observed reduction in performance might be due to the models’ reliance on reading frame. However, these datasets are also out-of-distribution in other ways, so this is not a definitive test.

### A.1 Accuracy Metric

We use a metric of top-kl accuracy, a generalization of top-k accuracy, which is defined as such: if there are *k* positives in the true labels, take the top *kl* predictions made by the model. Then compute the fraction of true positives covered. As *l* increases, top-kl increases (non-strictly) monotonically. When *l* = 1, this is equivalent to top-k accuracy, the value of recall and precision at the point where the two quantities are equal.

We provide top-kl scores for a variety of *l* values between 1 and 100, to demonstrate the degree to which these models are not correctly identifying splice sites in these settings. See Figure 6a for an illustration of this metric.

**Figure 6:**
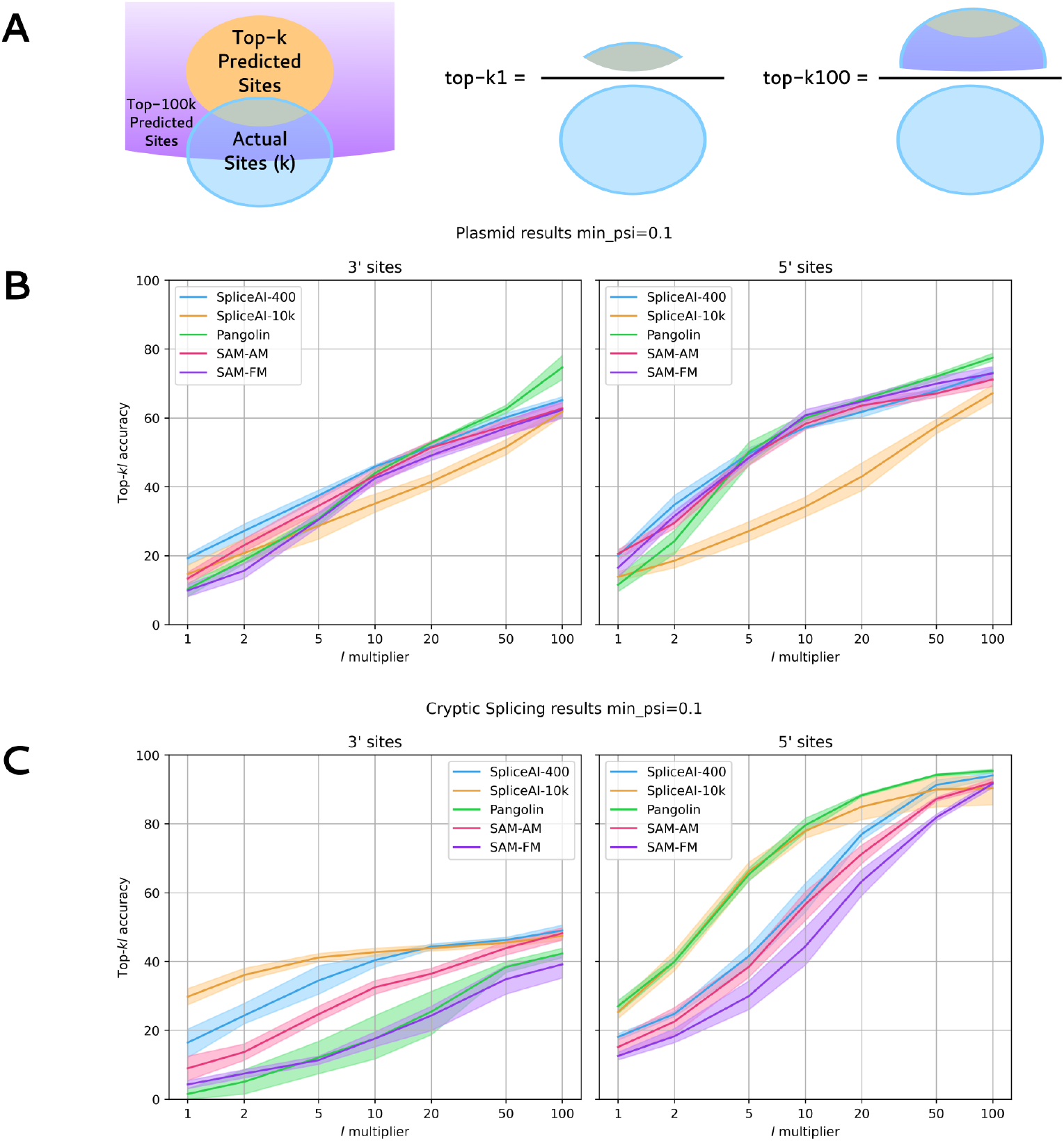
Evaluation of models on non-genomic datasets. (a) Illustration of top-kl accuracy. (b) Plasmid Results. (c) Cryptic Splicing Results.

### A.2 Plasmid Data

We use data from Anderson et al. [2024] in which plasmids (circular DNA constructs introduced into cultured cells) were used to explore the effects of repetitive CAG sequences on splicing. We use this splicing data, focusing on unique reads for our analysis, to compute top-kl scores by various models. As seen in Figure 6b, we find that all models generally do poorly on this task, only able to achieve ~50% coverage of true splice sites when predicting 10x as many as exist. Notably, one of the best genomic predictors, SpliceAI-10k, does particularly poorly on this task, while more mechanistic SAM models tend to do better for low *l*.

### A.3 Cryptic Splicing Data

This is based on a reanalysis of the RNA sequencing data from Kowal et al. [2025]. In this study, the body of an intron inserted into the 5’ untranslated region (UTR) of a green fluorescent protein (GFP) minigene reporter was randomized, and splicing at cryptic sites (sites not normally spliced) within the intron was assessed. We evaluate all models on these within-intron sequences. As seen in Figure 6c, results are similar to plasmids, though with a gap between 3SS and 5SS prediction accuracy, with 3SS being predicted with substantially lower accuracies.

## B Details on Triplet Replacement Experiments

In Figure 7, points are a mean over 5 model replicates; error bars indicate bootstrap confidence intervals over those replicates. Spikes below 0 indicate a drop in accuracy created by replacement by the codon, indicating that the model predicts that this codon suppresses the exon. Clear spikes for stop codons TAA, TAG, and TGA can be seen on SpliceAI and Pangolin models; these spikes are also largely phase-specific, only taking place for Phase 0 (in-frame) codons. For the SAM-FM and SAM-AM models, this effect only appears for TAG and TAA codons, and not the TGA codon. Additionally, there is no strong phase-specific effect.

**Figure 7:**
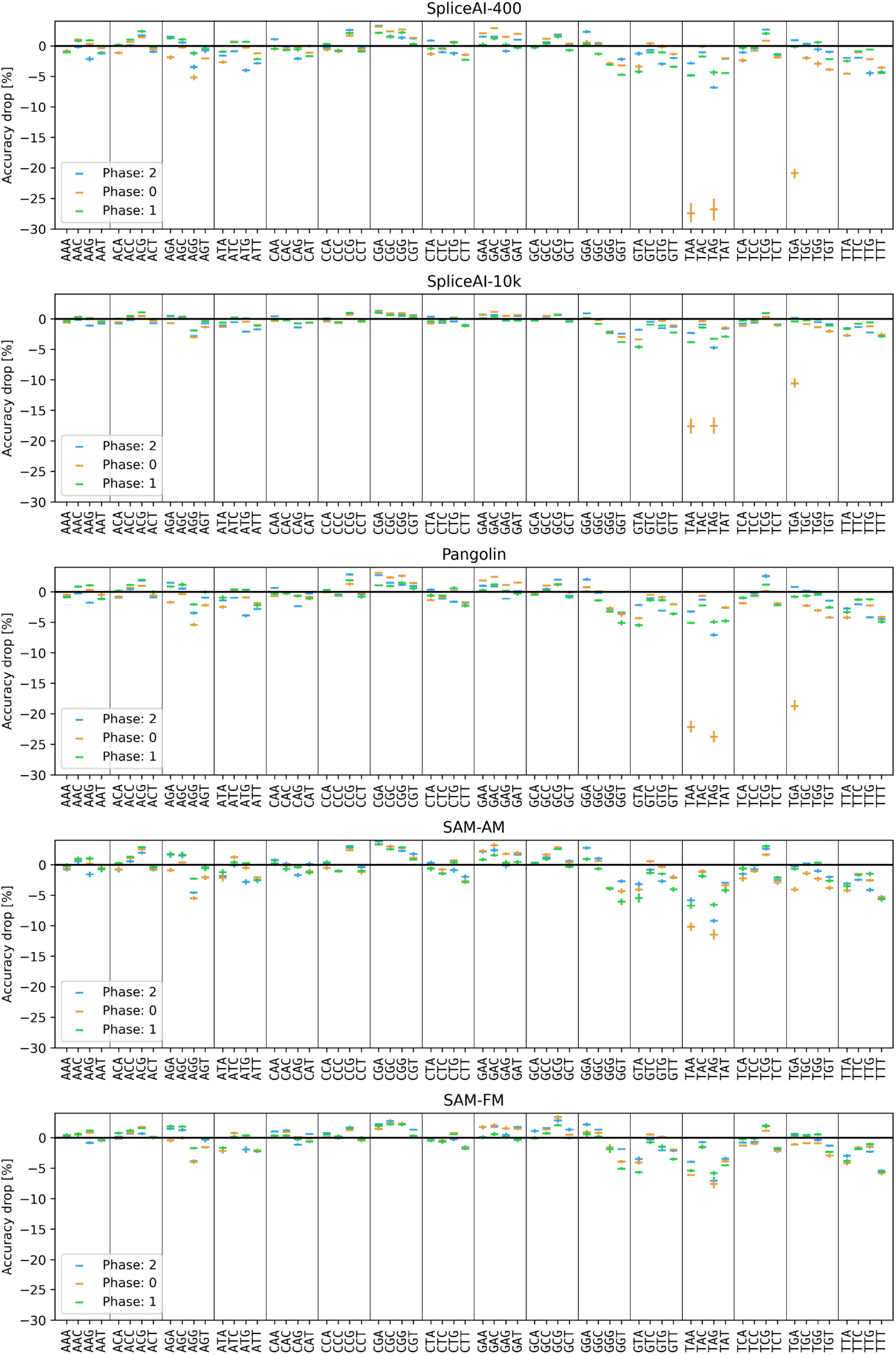
Individual triplet/phase results for the in-silico replacement experiments, one graph per model.

## C Is there a Global ORF?

Broadly, we find that the models are not learning a global ORF, but rather are treating each exon individually. Below are two experiments to this effect.

### C.1 Deletion Effect on Neighboring Exons

In this experiment, we analyze the effects of deletions in one particular exon on the nearby splice sites in adjacent exons. The results in Figure 8 indicate that the models are learning to recognize the frame coherence of a single exon but are not recognizing that an entire transcript needs to have a frame coherent coding region. Either they are incapable of this long-range computation, or the need to produce marginal probabilities of individual splice sites makes such information difficult to use in adjusting splice site scores.

**Figure 8:**
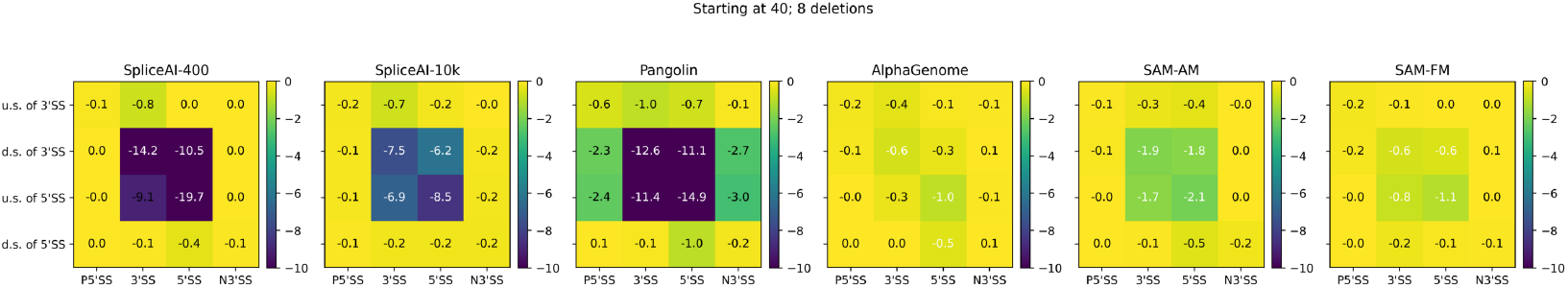
Per-model deletion effect on neighboring exons. Each panel is a separate model, with the y-axis being the deletion location and the x-axis being the affected splice site. The color indicates the change in prediction accuracy for the splice site. For all models except Pangolin, there is no effect, and for Pangolin the effect is 5 times smaller than the effects on the current exon.

### C.2 Two Exon Frame Repair Experiment

We more deeply investigate the effects of frame-disrupting mutations on exons other than the one the deletion takes place in. This experiment is similar to the deletion experiment explored in Figure 2,but using deletions 10nt away from the splice sites to decrease the effect on the exon in which the deletion is occurring.

As visualized in Figure 9a, we investigate deletions of X=1 or 2 in the first exon and Y=2 or 1 in the second exon, each of which disrupts the frame by itself, but whose effects on the overall frame “cancel out”, leaving the first and second exon mostly in the same frame.

**Figure 9:**
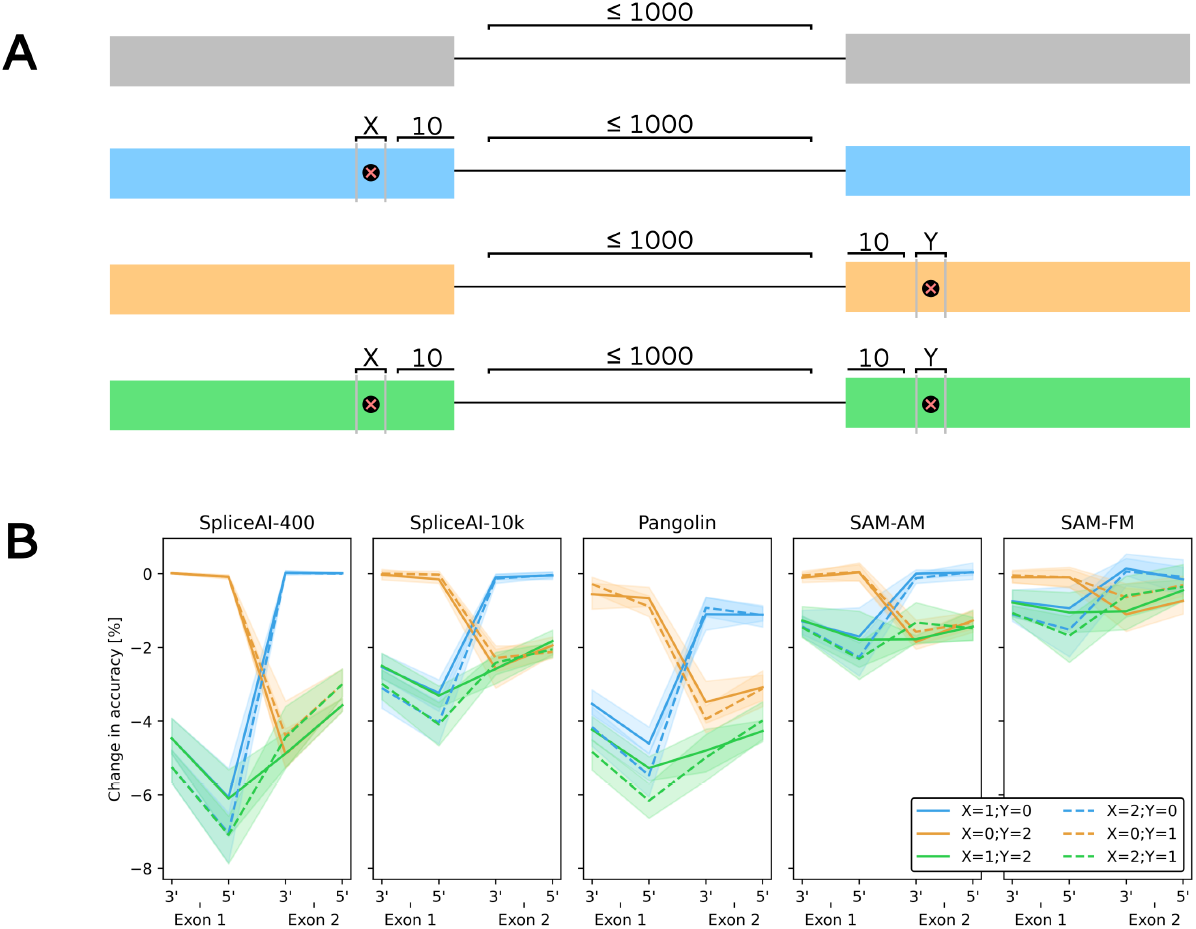
Two Exon Frame Repair Experiment. (a) Experiment Description. X and Y are the deletions in each exon. (b) Results of the Two Exon Frame Repair Experiment. The green conditions have complementary deletions in each exon, which should repair the frame, while the blue and orange conditions do not, and so have unrepaired frames.

If the models’ use of frame alignment information is within-exon, as we theorize, we should see that the effects of the X deletion and Y deletion are effectively independent of each other, with the accuracies superposing. However, if the models are using multiple-exon information, we should find that making these two complementary deletions leads to less of an accuracy impact than the sum of their individual effects.

As can be seen in Figure 9b, for all models, the effects are additive, as we expect. We can thus conclude that none of the models we investigate are identifying a coherent frame across multiple exons. Blue lines represent deletions in only the first exon, orange lines only in the second, and green in both; dashes are used to distinguish the two ways each of these conditions can be accomplished.

## D Robustness of the FPA Metric

### D.1 Alternative Metric: Fraction Actual Below

We “transpose” the “fraction of predictions above” metric described in Figure 4. In FPA, we look at control elements with a similar actual splicing level, and then see what percentage of them have a higher predicted score.

In our alternative metric, fraction actual below, we look at control elements with a similar predicted splicing score, and then see what percentage of them have a lower actual splicing value. These are roughly equivalent metrics. We find (Figure 10) that results are directionally similar, with the main differences being that SAM models score lower under both minigene datasets on this metric. This does not change any qualitative results.

**Figure 10:**
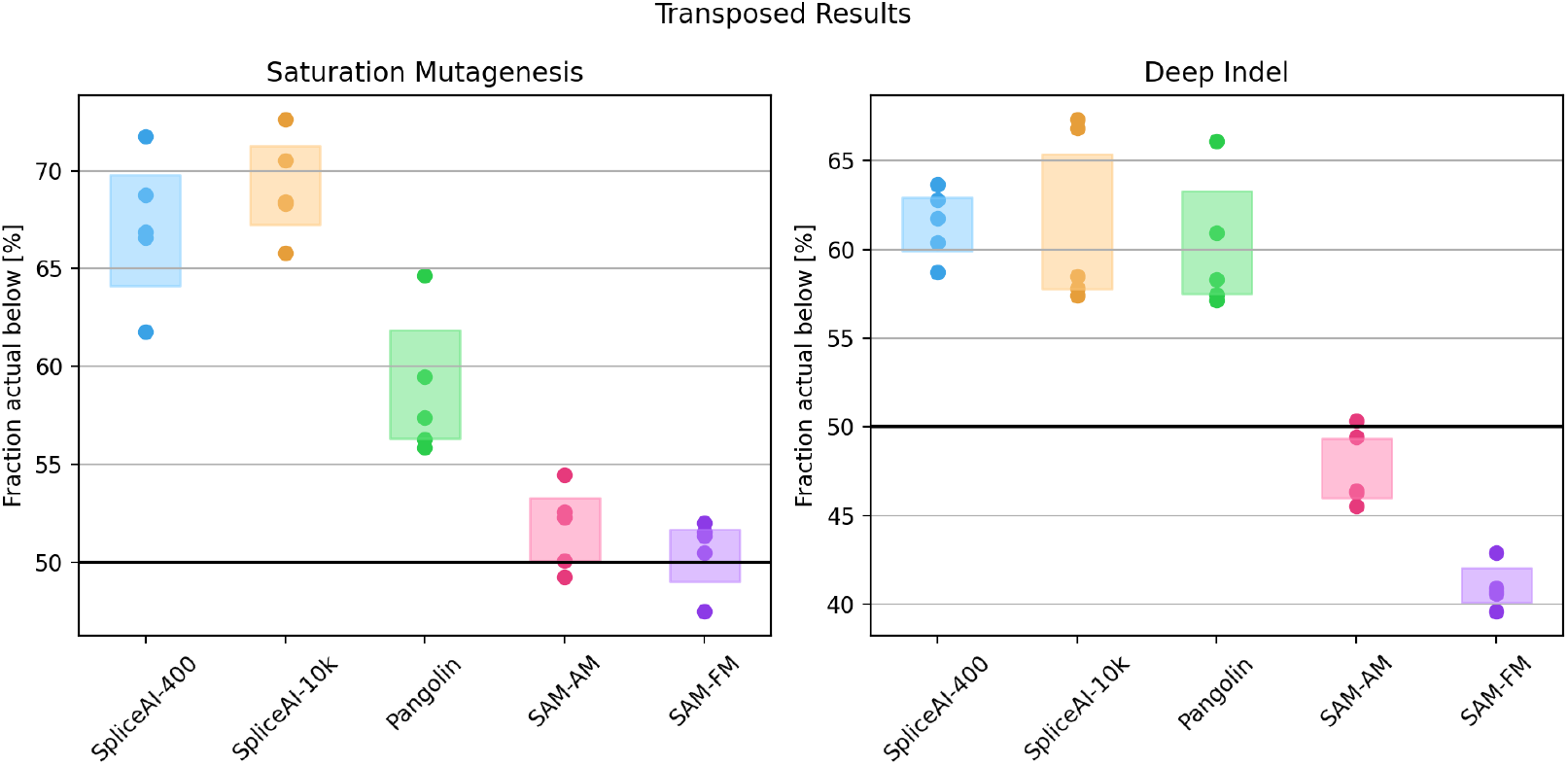
Fraction Actual Below metric for the minigene experiments.

### D.2 Sensitivity to the Number of Control Exons

We vary the size of the control set between 33 and 300 exons (Figure 11) and see that the results are flat; Section 4’s results do not depend on the particular value chosen.

**Figure 11:**
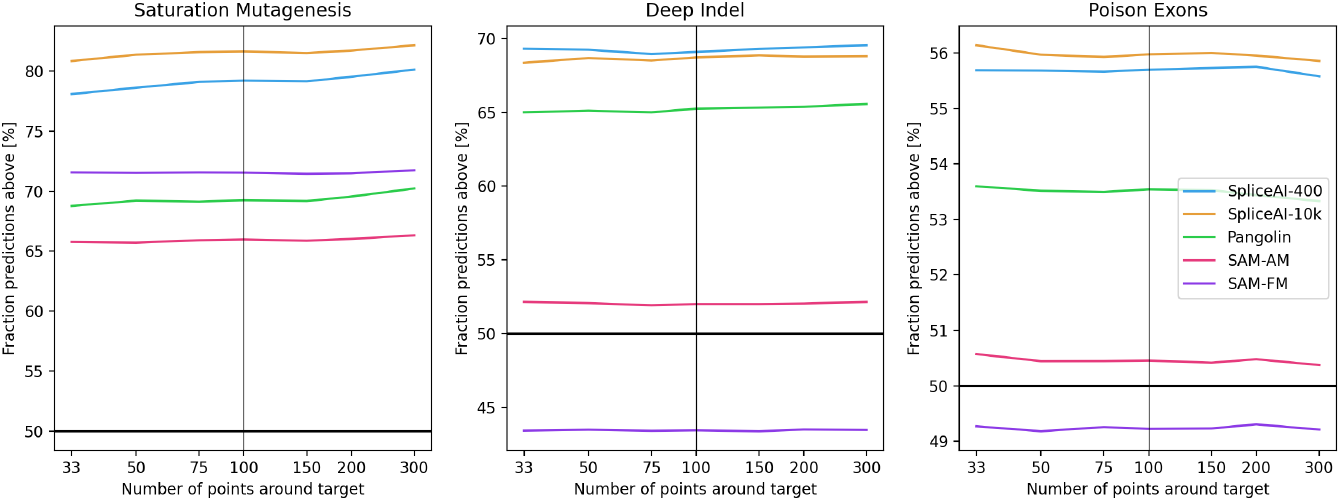
Fraction of predictions above as a function of the number of control exons used per target exon, for each of the three datasets. The vertical line marks the value of 100 used in the main paper.

**Figure 12:**
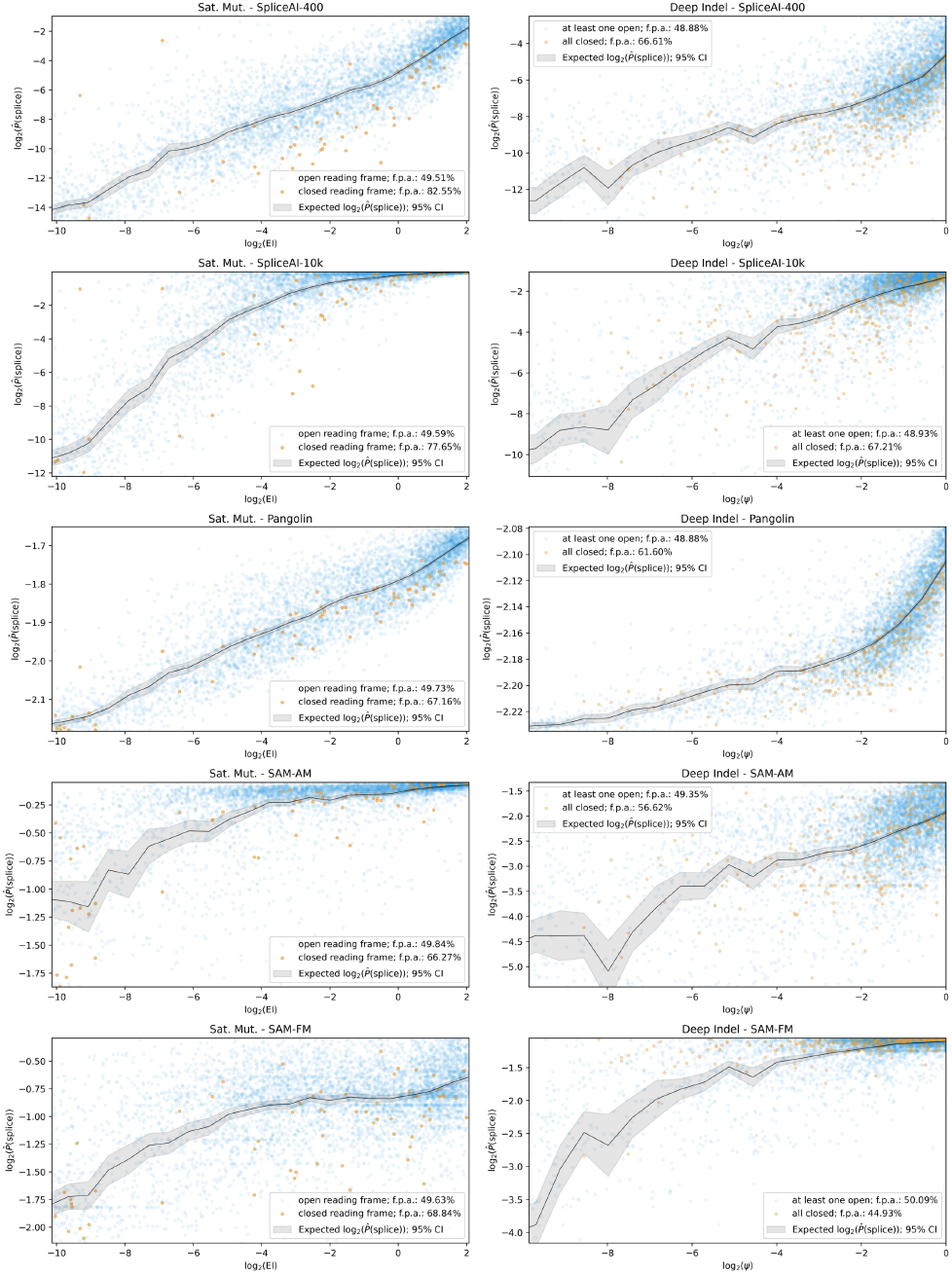
Scatterplots for the minigene experiments, for one replicate. Left: Saturation Mutagenesis results. Right: Deep Indel results (*ψ* denotes PSI). Legends give the FPA (Figure 4) of each group of exons.

## Scatterplots for the real data experiments

For Saturation Mutagenesis, the plot is log_2_(EI) on the x axis plotted against log_2_(predicted splice value) on the y axis; for Deep Indel the x axis is instead log_2_(PSI). The black line is a binned prediction of the log predicted splice value based on 25 bins of the measured value. We then plot samples with an open reading frame (blue) and closed reading frame (orange). As seen in Figure 12, in Saturation Mutagenesis, we find that on closed reading frames, the models underestimate the EI, and on Deep Indel we find the same phenomenon for the SpliceAI and Pangolin models, but a slight overestimation of the target splice site values for SAM-FM, and little bias for SAM-AM.

In Figure 13 we plot the raw results of the poison exon experiment for the first replicate on each model in a scatterplot, with a linear regression plotted as a line. We find that in general there is a lot of error in these predictions, but the overall predictions are always positively correlated with EF 5%. Additionally, we find that for SpliceAI and Pangolin models, the “Closed” line (all 3 frames closed) is generally lower than the “Open” line, indicating a depression in prediction of the exon, even after controlling for EF 5%. A more rigorous analysis of this can be found in Section 4.

**Figure 13:**
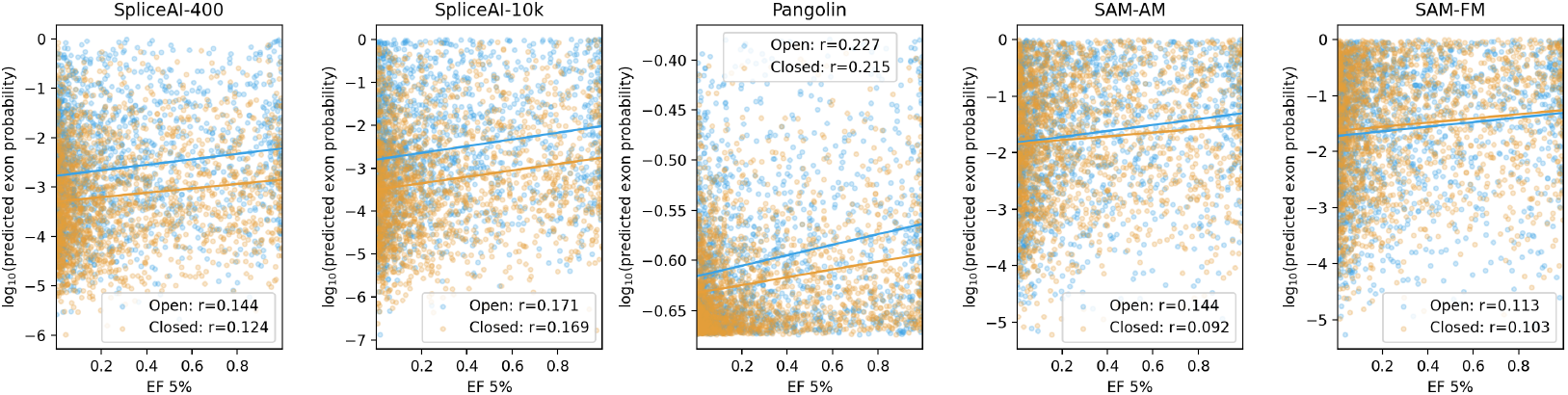
Full results of Poison Exon Experiment for one replicate.

## Footnotes

2 Throughout this paper we use T rather than U, even when discussing RNA.

3 In some cases we “substitute” a triplet with itself; this should not have a large effect on aggregate statistics.

4 We exclude from this analysis any replacements that introduce or remove a stop codon in a frame other than the one of the 3mer being applied, e.g., **GGT**AG *→* **CCC**AG and its reverse are excluded.

5 The average of splice site scores.

6 EI, or “exon inclusion”, and PSI, or “percent spliced in”, are the two studies’ measures of the level at which an exon is included in the final mRNA versus being spliced out as part of the surrounding intron.

